# The primary cilium is required for MC4R control of food intake and body weight

**DOI:** 10.1101/2020.11.13.382234

**Authors:** Yi Wang, Adelaide Bernard, Fanny Comblain, Xinyu Yue, Christophe Paillart, Sumei Zhang, Jeremy F. Reiter, Christian Vaisse

**Affiliations:** Department of Medicine and The Diabetes Center, University of California, San Francisco, California 94143-0540 USA; Department of Biochemistry and Biophysics, Cardiovascular Research Institute, University of California, San Francisco, California 94158-2324 USA; Chan Zuckerberg Biohub, San Francisco, CA 94158, USA

**Author notes:** Y.Wang, A.Bernard & F.Comblain contributed equally to these studies. Correspondence should be addressed to C.V.

## Abstract

The Melanocortin-4 Receptor (MC4R) plays a critical role in the long-term regulation of energy homeostasis and mutations in MC4R are the most common cause of monogenic obesity. However, the precise molecular and cellular mechanisms underlying the maintenance of energy balance within MC4R expressing neurons are unknown. We recently reported that MC4R localizes to primary cilia, a cellular organelle that allows for partitioning of incoming cellular signals, raising the question of whether MC4R functions there. Here, using mouse genetic approaches, we found that cilia are required specifically on MC4R-expressing neurons to restrain feeding behavior. Moreover, these cilia were critical for pharmacological activators of MC4R to exert an anorexigenic effect. MC4R is expressed in multiple brain regions. Using targeted deletion of primary cilia, we found that cilia in the paraventricular nucleus (PVN) of the hypothalamus are essential to restrict food intake. MC4R activation increases adenylyl cyclase activity. Like removing cilia, inhibiting adenylyl cyclase activity in the cilia of MC4R-expressing neurons of the PVN caused hyperphagia and obesity. Thus, MC4R signals via cilia of PVN neurons to control food intake and body weight. We propose that defects in ciliary localization of MC4R cause obesity in human inherited obesity syndromes and ciliopathies.

## INTRODUCTION

Most mammalian cells, including neurons, possess a single, immotile primary cilium, an organelle that transduces select signals^1^. Defects in the genesis or function of primary cilia cause a range of overlapping human diseases, collectively termed ciliopathies^2^. Several ciliopathies, such as Bardet-Biedl syndrome and Alström syndrome, cause obesity, and mutations in genes encoding ciliary proteins, such as *CEP19, ANKRD26* and *ADCY3*, cause non-syndromic obesity in mice and humans^3–6^. While the mechanisms underlying a number of cilia-associated phenotypes, such as polycystic kidney disease or retinal degeneration, have been at least partly elucidated, how ciliary dysfunction leads to obesity remains poorly understood^7,8^. Since primary cilia are essential for embryonic development, in particular through their critical role in Hedgehog signaling^9,10^, one hypothesis is that obesity could result from perturbations in the development of cells and pathways implicated in the regulation of energy homeostasis. However, ubiquitous ablation of the primary cilia in adult mice or from neurons also leads to obesity^11^. This finding indicates that neuronal primary cilia are post-developmentally required for the function of one or more signaling pathway implicated in the regulation of energy homeostasis.

We recently demonstrated that the melanocortin-4 receptor (MC4R) localizes to the neuronal primary cilia *in vivo*^12^. MC4R is a G protein-coupled receptor (GPCR) essential for long-term regulation of energy homeostasis^13^. In humans, polymorphisms at the *MC4R* locus are tightly associated with obesity in genome wide association studies^14^, heterozygous mutations in the *MC4R* coding sequence are the most common monogenic cause of severe obesity, and individuals with homozygous null mutations display severe, early-onset obesity^15–17^. In mice, deletion of *Mc4r* causes hyperphagia and severe obesity^18^. MC4R is found in a number of neuronal populations but its expression in the para-ventricular nucleus of the hypothalamus (PVN) is both necessary and sufficient for regulation of food intake and body weight^19^.

MC4R activity is regulated by two endogenous ligands, an anorexigenic agonist, alpha melanocyte-stimulating hormone (α-MSH), and an orexigenic antagonist/inverse agonist, Agouti-Related Peptide (AgRP). These neuropeptides are produced by neurons of the arcuate nucleus of the hypothalamus (ARC) under the control of the adipocyte-secreted hormone leptin^13^. As leptin levels are proportional to fat mass^20^, the observation that MC4R can localize to neuronal primary cilia suggests that signaling from the primary cilia of MC4R expressing neurons is a rate-limiting step in the sensing of energy stores and conditional modulation of energy intake.

Here, we use a combination of genetic and pharmacological approaches to demonstrate that primary cilia are required for ligand dependent activation of MC4R and long-term regulation of energy homeostasis by MC4R-expressing PVN neurons, and that altered MC4R ciliary function can underlie ciliopathy-associated obesity.

## RESULTS

### Genetic ablation of primary cilia in MC4R-expressing neurons phenocopies MC4R deficiency

We determined whether MC4R-expressing neurons require primary cilia to regulate body weight. Primary cilia can be specifically eliminated without affecting cell viability by inactivating *Ift88*, a gene encoding an intra-flagellar transport protein specifically required for ciliogenesis and ciliary maintenance^21^. To inactivate *Ift88* in MC4R neurons, we first generated an *Mc4r^t2aCre^* allele by CRISPR/Cas9-mediated zygotic recombination (Figure 1A). We inserted a T2A sequence and Cre recombinase open reading frame at the terminator codon of *Mc4r* so that the endogenous promoter/enhancer elements direct *Cre* expression to MC4R-expressing cells. Accurate insertion at the *MC4R* locus was verified by long-range PCR and sequencing. Recombinase activity, assessed in *Mc4r^t2aCre/t2aCre^ Rosa26^Ai14/Ai14^* mice which express a red fluorescent protein in a Cre-dependent fashion, recapitulated the endogenous MC4R expression in particular in the PVN^22^ (Figure 1N). Importantly, the weight curves of mice homozygous for the insertion (*Mc4r^t2aCre/t2aCre^*) did not differ from those of wild type littermates, indicating that the *Mc4r^t2aCre^* allele preserves MC4R function (Figure 1B).

**Figure 1.**
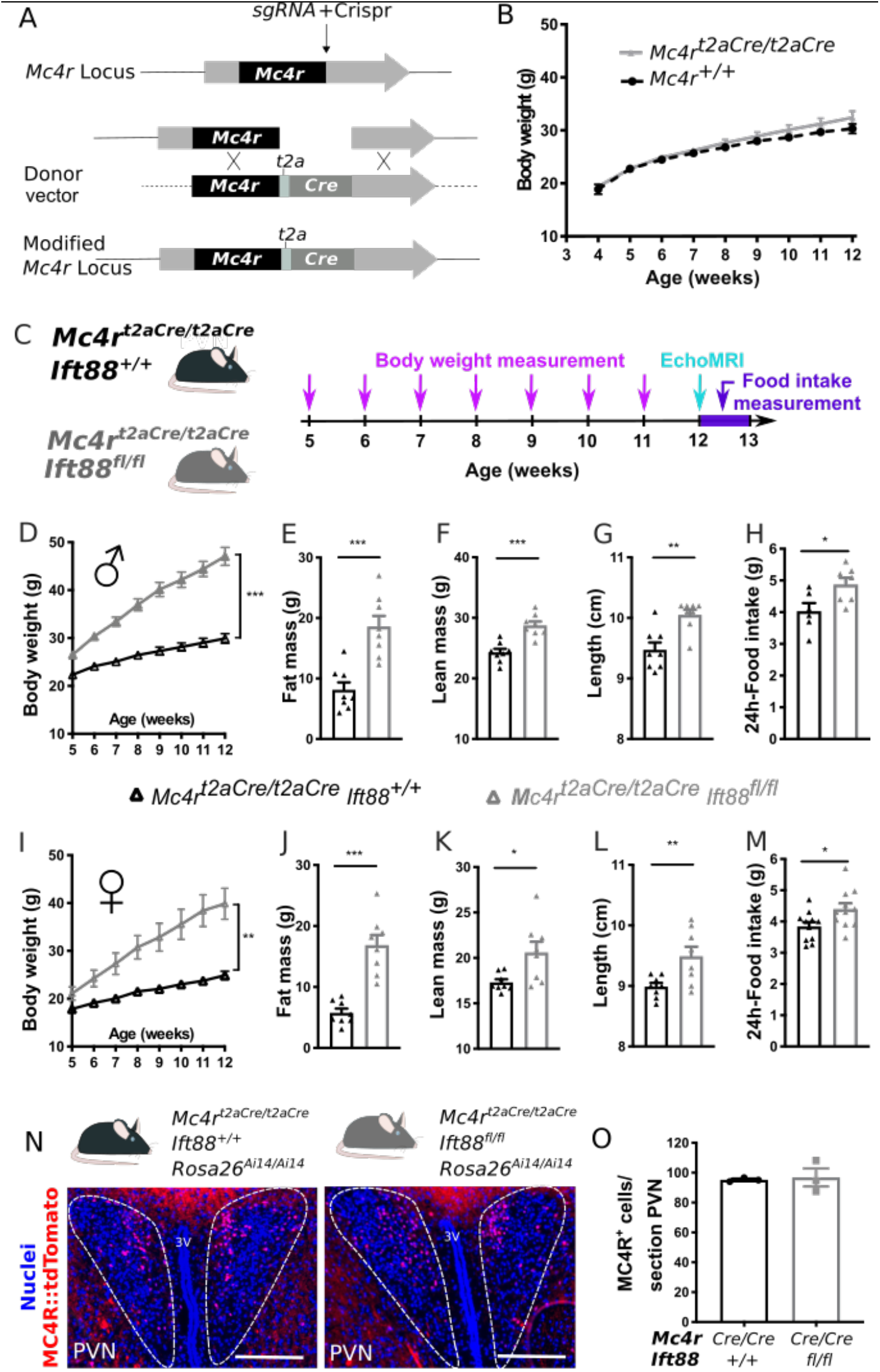
Deletion of *Ift88* in MC4R-expressing neurons leads to obesity. **A)** Strategy used to target the mouse *Mc4r* locus by CRISPR/Cas9 to insert a *T2A* and *Cre* open reading frame in frame with *MC4R*. **B)** Body weights of *Mc4r^t2aCre/t2aCre^* (n=7) mice and their *Mc4r^+/+^* littermates (n=6). **C)** Experimental design for phenotyping of *Mc4R^t2aCre/t2aCre^ Ift88^+/+^* and *Mc4R^t2aCre/t2aCre^ Ift88^fl/fl^* mice. **(D-M)** Phenotyping of control *Mc4R^t2aCre/t2aCre^ Ift88^+/+^* and *Mc4R^t2aCre/t2aCre^ Ift88^fl/fl^* male (D-H) and female (I-M) mice. Body weights were measured weekly from 5 to 12 weeks of age **(D,I)**. Fat mass **(E,J)** and lean mass **(F,K)** (measured by echoMRI), and length **(G,L)** were assessed at 12 weeks of age. n=8 mice per group. **H,M)** 24-hour food intake at 12 weeks of age (females n=11 control and 11 experimental; J, males n=6 control and 7 experimental). **N)** Representative images of the PVN in which MC4R-expressing neurons express a red fluorescent protein (MC4R::tdTomato) in both *Mc4r^t2aCre/t2aCre^ Ift88^+/+^* (left) and *Mc4r^t2aCre/t2aCre^ Ift88^fl/fl^* (right) mice, Scale bar, 200 μm. **O)** Quantification of the number of MC4R-expressing neurons in *Mc4r^t2aCre/t2aCre^ Ift88^+/+^* and *Mc4r^t2aCre/t2aCre^ Ift88^fl/fl^* mice (n=3 PVN sections per group, ns). All values are displayed as mean ± SEM. *p<0.05, **p<0.01 ***p<0.001.

We crossed *Mc4r^t2aCre^* mice to *Ift88^fl/fl^* mice^21^ to generate mice with specific deletion of *Ift88* in MC4R-expressing cells. To determine the effect of primary cilia deletion in MC4R neurons, we compared *Mc4r^t2aCre/t2aCre^ Ift88^fl/fl^* mice to *Mc4r^t2aCre/t2aCre^ Ift88^+/+^* littermates (Figure 1C). *Mc4r^t2aCre/t2aCre^ Ift88^fl/fl^* mice were born at a Mendelian ratio and develop a severe obesity phenotype (Figure 1D-M) that mimics germline loss of MC4R^18^. Specifically, body weight curves of *Mc4r^t2aCre/t2aCre^ Ift88^fl/fl^* and *Mc4r^t2aCre/t2aCre^ Ift88^+/+^* mice diverged after weaning (Figure 1D,I). This difference in body weight was characterized by a large increase in fat mass (Figure 1E,J), an increase in lean mass (Figure 1F,K) and length (Figure 1G,L) as well as an increase in food intake (Figure 1H,M). We observed the same effect in males (Figure 1D-H) and females (Figure (1I-M).

To confirm that loss of primary cilia in MC4R neurons did not affect neuronal survival, we generated triple knock-in *Mc4r^t2aCre/t2aCre^ Ift88^fl/fl^ Rosa26^Ai14/Ai14^* mice. We found no difference in the number of MC4R-expressing neurons in the PVNs of *Mc4r^t2aCre/t2aCre^ Ift88^fl/fl^ Rosa26^Ai14/Ai14^* mice when compared to those of *Mc4r^t2aCre/t2aCre^ Ift88^+/+^ Rosa26^Ai14/Ai14^* control mice (Figure 1N,O). Together these results indicate that primary cilia are critically and cell autonomously required for the function of MC4R neurons.

### Primary cilia are required for anorexigenic MC4R signaling

Since loss of primary cilia does not appear to affect the survival of MC4R-expressing neurons (Figure 1), we hypothesized that the obesity observed in *Mc4r^t2aCre/t2aCre^ Ift88^fl/fl^* mice was due to impaired MC4R function in differentiated neurons. To test this hypothesis, we determined whether primary cilia are required for the anorexigenic agonist-dependent function of MC4R in adult mice.

As opposed to ubiquitous germline ablation of primary cilia, which is embryonic lethal, ubiquitous adult conditional ablation of primary cilia, achieved by deleting *Ift88* using a tamoxifen-inducible Cre (*UBC-cre/ERT2 Ift88^fl/fl^*), results in obesity^11^.

To determine whether loss of primary cilia affects MC4R signaling, we assessed whether the timeline of primary cilia ablation and onset of the metabolic phenotype in this model is compatible with a dysfunction of the central melanocortin system. Adult *UBC-cre/ERT2 Ift88^fl/fl^* mice injected with tamoxifen were compared to *Ift88^fl/fl^* mice injected with tamoxifen controls (Figure 2A). Tamoxifen led to neuronal ablation of the primary cilia specifically within *UBC-cre/ERT2 Ift88^fl/fl^* mice within two weeks, as assessed by ciliary ADCY3 staining, a delay consistent with the half-life of IFT88 (Supplementary Figure 1). Body weight, body composition, food intake and energy expenditure were assessed repeatedly over 4 weeks (Figure 2A,B). Concurrent with neuronal cilia loss, tamoxifen-injected *UBC-cre/ERT2 Ift88^fl/fl^* mice developed obesity characterized by increased fat mass and a slight increase in lean mass (Figure 2B-E). This accumulation of fat mass was associated with hyperphagia, rather than changes in energy expenditure (Figure 2F,G), which would be consistent with loss of MC4R function in the PVN^23^.

**Figure 2:**
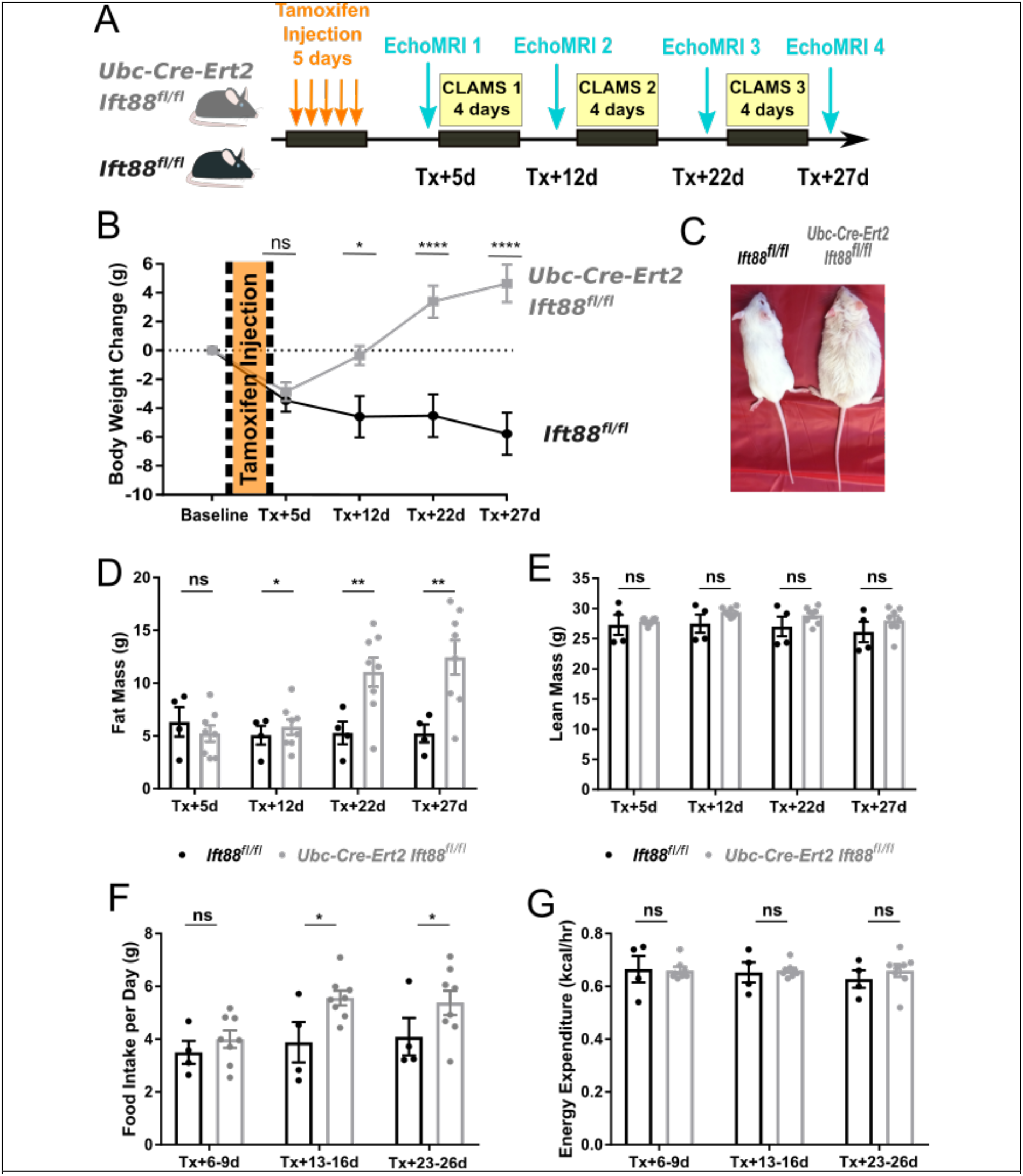
Time course of metabolic changes following ablation of primary cilia in adult mice. **A)** Schematic of the experimental protocol. **B)** Time course of body weight changes of *Ubc-Cre-Ert2 Ift88^fl/fl^* (n=8) and *Ift88^fl/fl^* (n=4) 20 week-old male mice at baseline and at the indicated times after tamoxifen (Tx) injection. **C)** Male control (*Ift88*^fl/fl^) and *Ubc-Cre-Ert2 Ift88^fl/fl^* mice 4 weeks following tamoxifen injection. **D)** Fat mass, **E)** lean mass as measured by EchoMRI, **F)** food intake and **G)** energy expenditure as measured in CLAMS, of *Ubc-Cre-Ert2 Ift88^fl/fl^* compared to *Ift88^fl/fl^* male mice at the indicated times after tamoxifen injection. Error bars represent SEM. *p < 0.05; **p < 0.01; ****p<0.0001.

To more specifically determine whether MC4R activity was impaired following ablation of primary cilia in this model, we evaluated the anorexigenic effect of pharmacological stimulation of MC4R before and after tamoxifen-induced ablation of primary cilia, but prior to weight divergence (Figure 3C). Specifically, *Ubc-Cre-Ert2 Ift88^fl/fl^ and control Ift88^fl/fl^* mice were injected intracerebroventricularly (ICV) with Melanotan-II (MTII), an MC4R agonist, before and after tamoxifen-mediated cilia loss (Fig. 3B). Mice were fasted for 24 hours prior to ICV injection, and their food intake was measured over a 4-hour period following injection. As expected, prior to tamoxifen-mediated cilia loss, MTII induced a drastic reduction of food intake in both groups compared to injection of the control solution, artificial cerebrospinal fluid (aCSF, Figure 3D). After tamoxifen injection, the anorexigenic effect of MTII gradually diminished specficially in *Ubc-Cre-Ert2 Ift88^fl/fl^* mice (Fig. 3D). Thus, primary cilia are necessary for MC4R-mediated effects on food intake.

**Figure 3:**
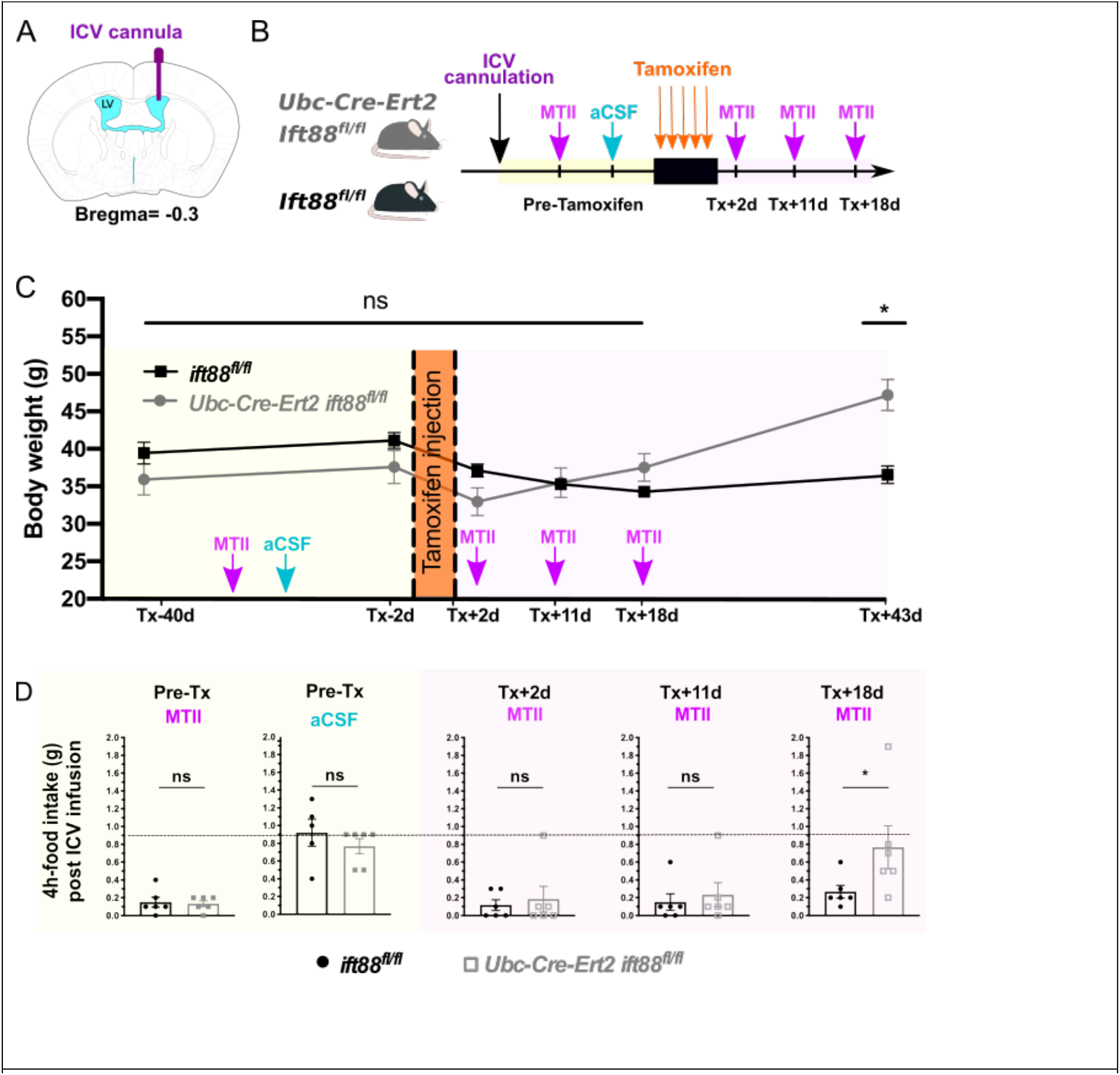
Primary cilia are essential for the response to the MC4R agonist MTII. **A)** Schematic of the placement of an ICV cannula in the lateral ventricle (coordinates: AP=-0.3, ML=+1, DV=-2.5) **B)** Schematic of the experimental protocol: over 20 week-old male control (*Ift88^fl/fl^*) and *Ubc-Cre-Ert2 Ift88^fl/fl^* littermates (n=6 per group) were implanted with an ICV cannula in the lateral ventricle. After recovery, food intake was measured following ICV delivery of MTII and vehicle control (aCSF). Mice were then injected with tamoxifen (Tx) for 5 days and the response to ICV-delivered MTII was measured again 2, 11, and 18 days after the last Tx injection. **C)** Weights of *Ift88^fl/fl^* control and *Ubc-Cre-Ert2 Ift88^fl/fl^* littermate mice during the experiment. **D)** Assessment of anorexigenic effect of MTII (0.5 nmol) on short-term food intake (4 hours) compared to vehicle (aCSF) before and after primary cilia loss. Error bars represent SEM. * indicates p<0.05.

As an additional test of whether cilia are required for MC4R signaling, we tested the effects of deleting cilia on the activity of THIQ, another specific agonist of MC4R that inhibits food intake. Similar to MTII, the anorexigenic response to ICV administration of THIQ was abolished specifically following removal of primary cilia specifically in *Ubc-Cre-Ert2 Ift88^fl/fl^* mice (Supplementary Figure 2).

As observed when specifically removing primary cilia from MC4R neurons developmentally, deletion of *Ift88* from adult hypothalamus did not alter neuron number (Supplementary Figure 3). Deletion of *Ift88* also did not alter expression levels of *Sim1* or *Mc4r* in the adult hypothalamus (Supplementary Figure 3D). These results indicate that the loss of responsiveness to MC4R stimulation upon removal of primary cilia was not due to loss of MC4R-expressing cells or decreased *Mc4r* expression.

In addition to regulating feeding behavior, PVN neurons respond to osmotic stimulation, for example, by phosphorylating the ribosomal protein S6^24^. pS6 activation by osmotic stimulation in PVN neurons was preserved in tamoxifen-treated *Ubc-Cre-Ert2 Ift88^fl/fl^* mice following cilia loss, indicating that primary cilia are not required for all functions of these neurons (Supplementary Figure 3E). Together, these data demonstrate that primary cilia are required in PVN neurons for MC4R regulation of food intake.

### Adult PVN primary cilia are required for anorexigenic MC4R signaling

MC4R is expressed in a number of brain regions, including multiple hypothalamic nuclei^25^. However, much of the anorexigenic activity of MC4R is due to its function in PVN neurons of the hypothalamus, where MC4R activity is both necessary and sufficient to inhibit food intake and control body weight^19,23^. In *Mc4r^t2aCre/t2aCre^ Ift88^fl/fl^* mice, primary cilia were ablated from all MC4R expressing cells, including those of the PVN. In *Ubc-Cre-Ert2 Ift88^fl/fl^* mice, treatment with tamoxifen removed cilia globally. Therefore, to test whether primary cilia are specifically required in the PVN for regulation of body weight through MC4R activation, we deleted *Ift88* by bilateral stereotaxic injection of adeno-associated virus (AAV) expressing GFP-tagged Cre recombinase (AAV-creGFP) into the PVN of adult *Ift88^fl/fl^* mice (Fig. 4A). Mice injected with adeno-associated virus expressing a GFP-tagged functionally impaired Cre recombinase (AAV-nGFP) served as controls. GFP expression indicated infected cells and allowed for post-hoc confirmation of PVN injection (Fig. 4C, D). Injection of Cre-producing virus, but not control virus, lead to the ablation of primary cilia in the PVN (Fig. 4C,D inserts).

**Figure 4:**
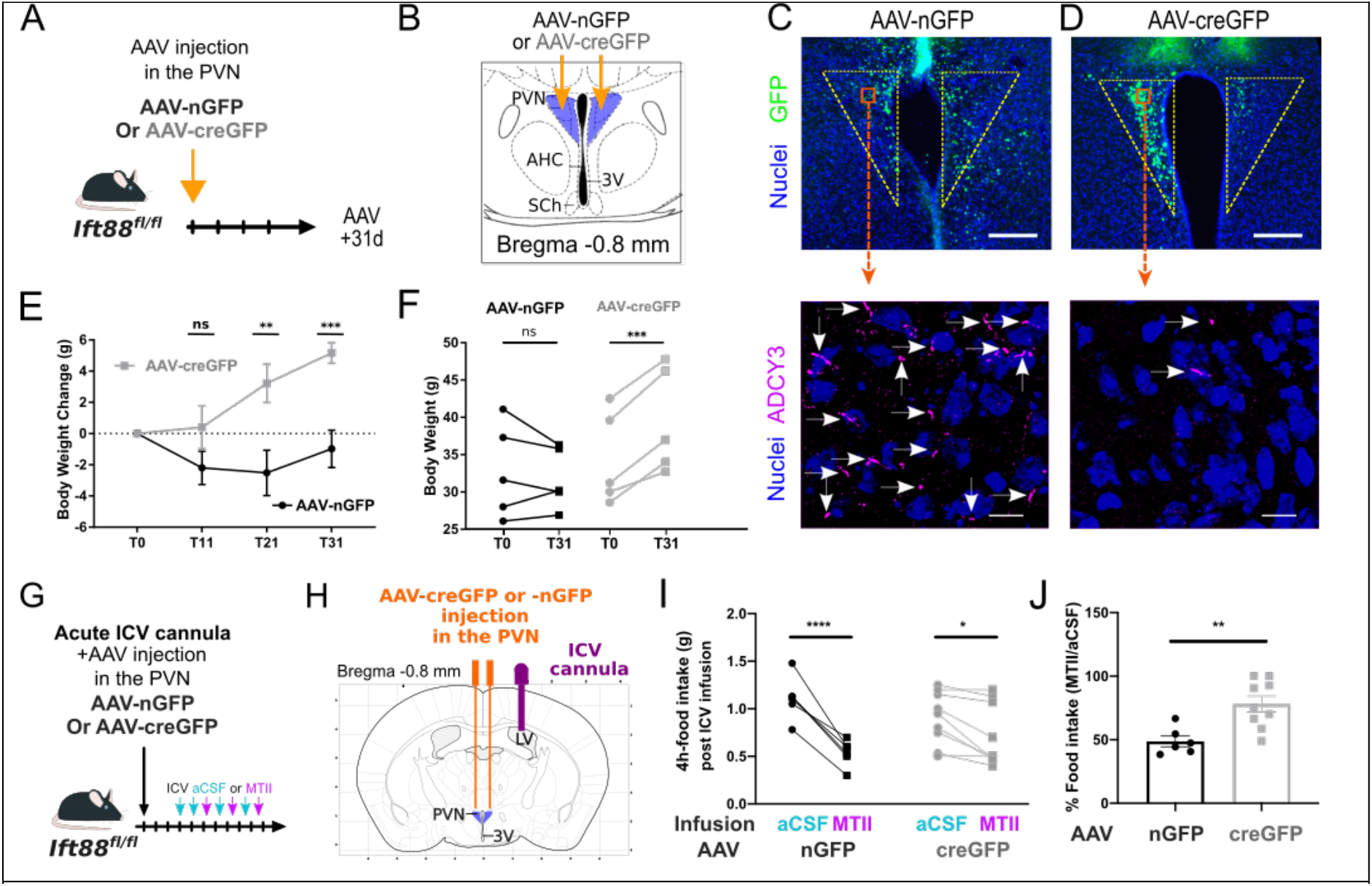
Primary cilia are required in PVN neurons for weight control and sensitivity to MC4R agonist. **A)** Schematic of the experimental protocol. Bilateral stereotaxic injections (coordinates: AP=-0.8, ML=±0.2, DV=-5.2) of AAV-creGFP or AAV-nGFP were performed on 20 week-old *Ift88*^fl/fl^ mice **B)** Schematic representation of hypothalamic region studied. **C,D)** Representative images of PVN sections of AAV-creGFP- or AAV-nGFP-injected mice showing AAV-infected cells in green, and nuclei in blue. Scale bar, 200 μm. Inserts: Immunofluorescent imaging of primary cilia (ADCY3, magenta) of PVN regions denoted in C and D. Arrows indicate cilia. Scale bar, 10μm. **E)** Body weights of *Ift88*^fl/fl^ mice following bilateral PVN injection of AAV-creGFP or AAV-nGFP (n=5 per group). **F)** Body weights at time of AAV injection and one month afterwards. Individual mice are indicated by lines. **G)** Schematic of the experimental protocol for testing the anorexigenic effects of the MC4R agonist MTII. 3 weeks following AAV injection and cannulation, 20 week-old *Ift88*^fl/fl^ mice were alternately treated with vehicle (aCSF) or 0.05 nmol Melanotan II (MTII) by ICV infusion after fasting for 24 hours. aCSF and MTII infusions were alternated with a 4-day interval between infusions. Food intake during 4h re-feeding periods following 24h fasting were then averaged for aCSF and MTII (values in I and J). **H)** Schematic of bilateral stereotaxic injections (coordinates: AP= −0.8, ML=±0.2, DV=-5.2) of AAV-creGFP (n=9) or AAV-nGFP (n=6), and placement of an ICV cannula in the lateral ventricle (coordinates: AP=-0.3, ML=+1, DV=-2.5). **I)** 4h food intake following injection of aCSF or MTII in AAV-creGFP- and control AAV-nGFP-injected *Ift88*^fl/fl^ mice (Repeated measures averaged). **J)** Percentage of food ingested in 4h following ICV administration of MTII normalized to administration of vehicle (aCSF). T=days following AAV injection. Error bars represent SEM. Statistics: t-test (J) or Repeated measures of two-way ANOVA followed by Sidak’s multiple comparisons test were performed. *p<0.05, **p<0.01, ***p<0.001, ****p<0.0001. PVN: paraventricular nucleus of the hypothalamus; 3V: third ventricle; SCh: suprachiasmatic nucleus; AHC: anterior hypothalamus centralis; ICV: intracerebroventricular; LV: Lateral Ventricle.

Following ablation of the primary cilia in the PVN, AAC-creGFP-injected *Ift88^fl/fl^* mice gained weight (Fig. 4E,F). Thus, cilia are critical for the regulation of energy homeostasis in the PVN and account, in full or in part, for the obesity observed after ubiquitous disruption of primary cilia in adult mice. As MC4R localizes to primary cilia of neurons in the PVN^12^, these data further suggest that MC4R could function at PVN cilia to control body weight.

To determine whether MC4R activation requires PVN primary cilia, we implanted an ICV cannula in the lateral ventricle at the time of AAV injection (Fig. 4G,H) and measured changes in the anorectic effect of MTII after PVN primary cilia loss as described above. Mice were injected with MTII or vehicle (aCSF) after a 24 hour fast, and their food intake was consequently measured over 4 hours. (Fig. 4I). While control AAV-nGFP-injected mice decreased their food intake by sixty percent on average after MTII injection as compared to after aCSF injection, this response was blunted in AAV-CreGFP-injected mice lacking primary cilia in the PVN (Fig. 4I,J). Together, these results demonstrate that the anorexigenic function of MC4R requires primary cilia in the PVN.

### Adenylyl cyclase signaling in the primary cilia of MC4R-expressing neurons in the adult PVN is essential for controlling food intake and body weight

We reasoned that if MC4R functions at the primary cilia of PVN neurons, then inhibiting MC4R signaling specifically in these primary cilia would increase food intake and cause obesity. Activation of MC4R stimulates Gas to increase adenylyl cyclase activity^26^. Inhibition of adenylyl cyclase specifically at the primary cilia can be achieved by expression of a constitutively active version of the cilium-localized, Gai-coupled receptor, GPR88 (GPR88(G283H) or GPR88*)^27^.

To target MC4R neurons of the PVN, we stereotaxically injected a Cre-dependent AAV encoding a FLAG-tagged version of GPR88* (AAV DIO Flag-GPR88*) into the PVNs of 20 week-old *Mc4r^t2aCre/t2aCre^* male mice. To verify injection accuracy, PVN transduction, and Cre activity, we coinjected AAV expressing mCherry in a Cre-dependent manner (AAV DIO-mCherry). Control mice included *Mc4r^t2aCre/t2aCre^* mice injected only with AAV DIO-mCherry as well as *Mc4r^+/+^* littermates injected with AAV DIO-GPR88* and AAV DIO-mCherry (to control for Cre-independent effects of AAV DIO-GPR88*). Body weight was measured weekly and body composition was assessed by Echo-MRI 3, 6 and 9 weeks post AAV injections. Food intake and energy expenditure were assessed at 3 weeks post AAV injections by CLAMS.

*Mc4r^t2aCre/t2aCre^* mice injected with AAV DIO-GPR88* into the PVNs increased their body weight (Figure 5 E,F) and fat mass (Figure 5H). Food intake was higher in AAV DIO-GPR88*-inected *Mc4r^t2aCre/t2aCre^* mice (Figure 5I), but energy expenditure was unaffected 3 weeks post AAV injection (Figure 5J). Thus, GPR88*-mediated obesity was attributable to hyperphagia.

**Figure 5.**
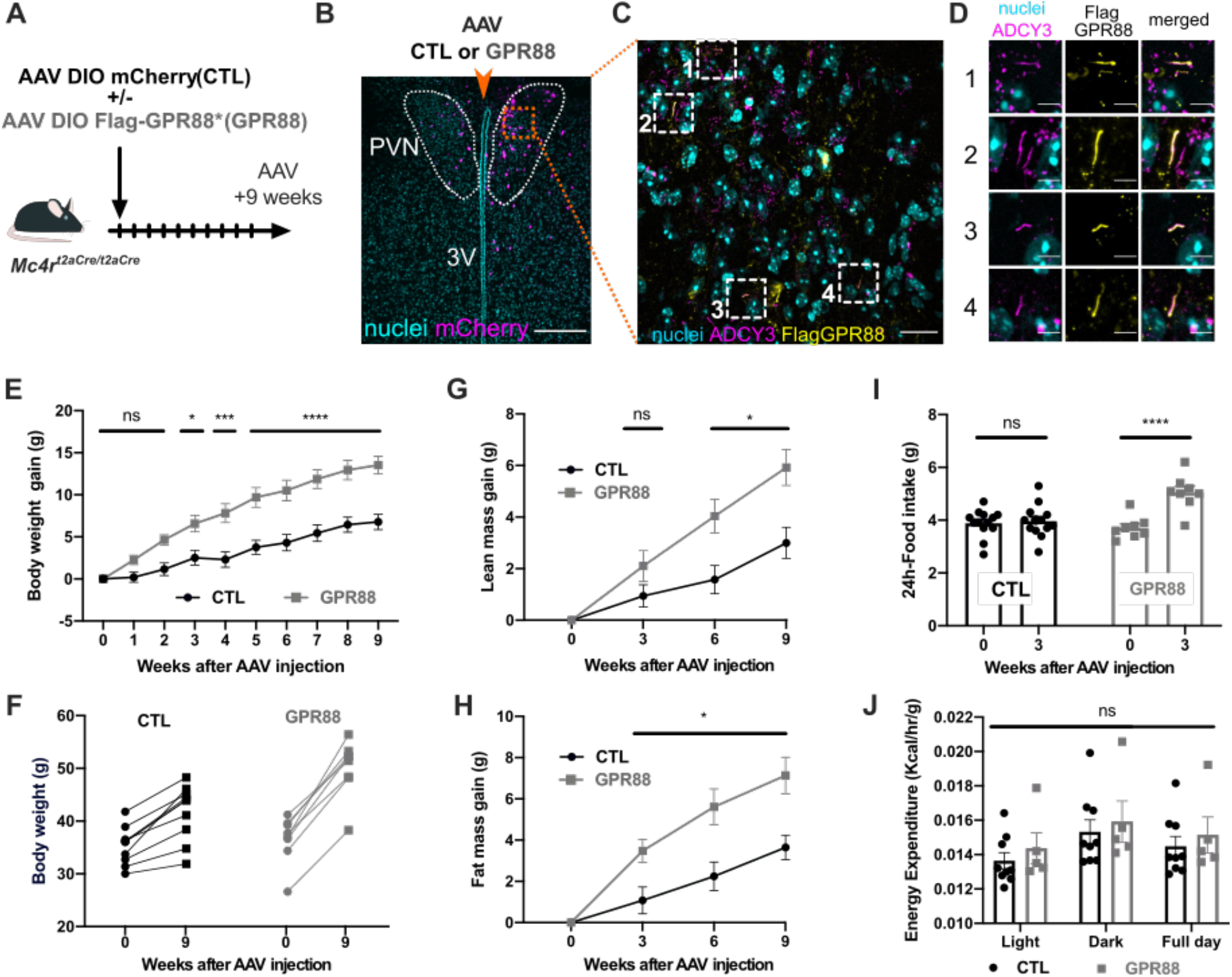
Inhibition of ciliary adenylyl cyclase causes hyperphagia and obesity. **A)** Schematic of the experimental protocol. Midline stereotaxic injections (coordinates: AP=-0.8, ML=0.0, DV=-5.5) of AAV DIO-FlagGPR88* and AAV DIO-mCherry in 18 weeks old *Mc4r^t2aCre/t2aCre^* male mice, hereinafter denoted as “GPR88”. “CTL” denotes *Mc4r^t2aCre/t2aCre^* mice injected with DIO-mCherry AAV only or wild type mice injected with AAV DIO-FlagGPR88*. **B)** PVN sections of GPR88 mice showing nuclei (cyan) and mCherry (magenta) indicating region of viral transfection. Scale bar, 200 μm. **C)** Inset from B. Immunofluoresence imaging of cilia (ADCY3, magenta), FLAG-GPR88* (yellow) and nuclei (cyan). Boxes indicate examples of localization of GPR88 to cilia of PVN neurons. Scale bar, 20 μm. **D)** Magnified insets from C depicting GPR88 localization to individual cilia. Scale bar, 5μm. **E)** Body weight change of GPR88 (n=8) and control (n=13) mice over the nine weeks following AAV injection. **F)** Weights at time of AAV injection and 9 weeks afterwards. Lines connect individual mice. **G)** Change in lean mass over the nine weeks following AAV injection. **H)** Change in fat mass over the nine weeks following AAV injection. **I)** 24h food intake at time of AAV injection and 3 weeks afterwards. **J)** Average hourly energy expenditure during light phase, dark phase and full day of GPR88 (n=5) and control (n=9) mice 3 weeks following AAV injection. All values are displayed as mean ± SEM. Statistics: Repeated measures of two-way ANOVA followed by Sidak’s multiple comparisons test were performed. *p<0.05, ***p<0.001 ****p<0.0001. 3V: third ventricle, PVN: paraventricular nucleus of the hypothalamus.

These results demonstrate that reducing adenylyl cyclase activity specifically at the primary cilia of adult MC4R PVN neurons is sufficient to increase food intake and disrupt regulation of body weight.

## DISCUSSION

Obesity is a hallmark of several human ciliopathies, including Bardet-Biedl syndrome, MORM syndrome and Carpenter syndrome. As the major physiological pathway implicated in the regulation of long-term energy homeostasis, the central leptin-melanocortin system may link human ciliopathies and obesity. Since deletion of primary cilia in adult mice causes leptin resistance, the initial hypothesis was that dysfunction of central leptin signaling could play a role in the observed associated obesity^11,28–30^. However, leptin resistance was subsequently found to be a consequence, rather than a cause, of obesity in this model^31^, suggesting that dysfunction of a downstream effector of leptin may account for ciliopathy-associated obesity.

Here, we used several complementary genetic and pharmacological approaches to demonstrate that MC4R, a GPCR critically required for maintenance of body weight, not only localizes to primary cilia, but also operates at primary cilia of PVN neurons to control energy homeostasis in adult mice. Removing primary cilia specifically from MC4R-expressing cells (in *Mc4r^t2aCre/t2aCre^ Ift88^fl/fl^* mice) phenocopies the obesity phenotype caused by germline loss of MC4R^18^, demonstrating that primary cilia are essential for the function of MC4R-expressing neurons.

As this requirement of primary cilia in MC4R-expressing neurons could reflect a developmental role for cilia caused by removing cilia, we investigated whether primary cilia are required for the production of MC4R-expressing neurons and found that removing cilia does not affect the number of MC4R-expressing neurons nor expression of MC4R. Therefore, primary cilia are dispensible for the production of MC4R-expressing neurons.

Moreover, three additional lines demonstrate that MC4R functions at primary cilia. First, cilia are essential for the anorexigenic effects of MC4R agonists such as MTII and THIQ. Second, cilia are required specifically in MC4R-expressing neurons for the regulation of feeding behavior. Third, inhibiting the MC4R signaling pathway by inhibiting adenylyl cyclase specifically in the primary cilia of MC4R expressing PVN neurons is sufficient to induce hyperphagia and obesity. Since obesity is the most striking phenotype resulting from ubiquitous ablation of primary cilia in adult mice, our data suggest that a major function of adult primary cilia is to regulate long-term energy homeostasis by transducing MC4R signaling in PVN neurons.

Since partial loss of MC4R activity, as caused by heterozygous *MC4R* mutations, is sufficient to cause severe obesity in mice and humans^17^, a parsimonious model for ciliopathy-associated obesity is that it results from decreased localization of MC4R to the primary cilia of PVN neurons. This model also implies that partial attenuation of the ciliary localization of MC4R is sufficient to cause obesity and that genetic variation decreasing primary cilia localization of MC4R could predispose to weight gain. It will be interesting to assess to what extent obesity-associated variants detected in GWAS studies are involved in primary ciliary function. In this respect, it is interesting to note that obesity-associated variants are located within the MC4R and ADCY3 region^32^ and that variants in the FTO region (the strongest obesity associated signal in GWAS) could affect the primary cilia transition zone component RPRGRIP1L^33^.

Coordinated feeding behavior results from adapting short-term feeding behavior to immediate food availability and long-term caloric needs to maintain body weight stable over time. While MC4R integrates neuro-endocrine signals provided by α-MSH and AgRP, MC4R-expressing neurons in the PVN are also sensitive to neural afferent inputs that communicate short-term energy status from arcuate nucleus glutamatergic neurons and GABAergic AgRP-expressing neurons^34,35^. Our findings suggest that primary cilia could be a mean for MC4R neurons to compartmentalize long-term energy state signaling resulting from MC4R activity modulation^13^. This subcellular compartmentalization of signaling may explain how MC4R neurons are able to integrate different timescales afferent information to coordinate feeding behavior. Future studies will address how MC4R neurons integrate ciliary and synaptic communication.

## EXPERIMENTAL PROCEDURES

### Origin of the mouse lines used

All animal procedures were approved by the Institutional Animal Care and Use Committee of the University of California, San Francisco. Mice were housed (with enrichment) in a barrier facility and maintained on a 12:12 light cycle (on: 0700-1900) at an ambient temperature of 23±2°C and relative humidity 50-70 %. Mice were fed with rodent diet 5058 (Lab Diet) and group-housed up to 5 or single housed after surgery. Experiments were performed with weight and sex-matched littermates. Mice expressing tdTomato in a Cre dependent manner, *Gt(ROSA)26Sor*^tm14(CAG-tdTomato)Hze^, and mice ubiquitously expressing Cre-Ert2, *(Tg(UBC-cre/ERT2)1Ejb, Ubc-Cre-Ert2)*, were obtained from Jackson Laboratories (Bar Harbor, ME). Mice carrying the *Ift88* conditional allele (*Ift88*^tm1.1Bky^, *Ift88^fl^*) were obtained from Bradley Yoder. Experimental male *Ubc-Cre-Ert2 Ift88^fl/fl^* and control *Ift88^fl/fl^* littermates on a mixed background were obtained by crossing *Ift88^fl/fl^* females with *Ubc-Cre-Ert2 Ift88^fl/fl^* males.

### Generation of *MC4R^t2aCre/t2aCre^* mice

Was performed at the Gladstone Institute mouse transgenic core as described previously for MC4R^egfp/egfp^ mice ^12^. Briefly, Super-ovulated female FVB/N mice (4 weeks old) were mated to FVB/N stud males. Fertilized zygotes were collected from oviducts and injected with (1) Cas9 protein (50 ng/ul), (2) a donor vector (20 ng/μl) consisting of 1kb of 5’flanking sequence (i.e. the MC4R coding sequence) followed by t2aCRE and 5.5 kb of 3’flanking sequence and (3) a sgRNA (25ng/μl) of which the guide sequence (GTCTAGCAGGTATTAAGTGGGGG) was designed to target nucleotides immediately downstream the MC4R stop codon in a short region that was not present in the donor vector into pronucleus. Injected zygotes were implanted into oviducts of pseudopregnant CD1 female mice. Pups were genotyped for insertions at the correct loci by PCR. MC4R neuron specific expression of Cre was verified by crossing mice with *Gt(ROSA)26Sor*^tm14(CAG-tdTomato)Hze^.

### Generation and injections of AAVs

AAV DJ-CMV GFP-Cre (AAV-creGFP) or AAV DJ-CMV GFP-ΔCre (AAV-nGFP), obtained from the Stanford Neuroscience Viral Core, were stereotaxically injected bilaterally (coordinates: AP=-0.8, ML=±0.2, DV=-5.5, volume= 2×0.5μL) in the brains of *Ift88*^fl/fl^ mice. Weight was measured for 1 month, after which mice were sacrificed to confirm the site of injection.

AAV DIO GPR88* plasmids were generated by replacing hChR2(H134R)-EYFP in pAA V-Ef1a-DIO-hChR2(H134R)-EYFP-WPRE-pA (obtained from K.Desseiroth, Stanford University) with GPR88(G283H)-FLAG. AAV DJ were prepared and titrated by the Stanford Neuroscience Viral Core which also provided the stock mCherry DIO-AAV (GVVC-AAV-14).

AAV DIO mCherry (0.2μL) +/- DIO GPR88 (0.8μL) were stereotaxically injected in the PVN (coordinates: AP= −0.8, ML=0.0, DV=-5.2) of *Mc4r^t2aCre/t2aCre^* mice to anatomically and genetically restrict expression to PVN MC4R expressing neurons. Controls were either wild-type mice injected with DIO-GPR88 + DIO-mCherry AAV or *Mc4r^t2aCre/t2aCre^* injected solely with AAV DIO-mCherry (0.2μL) diluted in artificial cerebro-spinal fluid (aCSF, 0.8μL).

### Stereotaxic Surgeries

Animals were anesthetized with an initial flow of 4% isoflurane, maintained under anesthesia using 2% isoflurane and kept at 30-37°C using a custom heating pad. The surgery was performed using aseptic and stereotaxic techniques. Briefly, the animals were put into a stereotaxic frame (KOPF Model 1900, USA), the scalp was opened, the planarity of the skull was adjusted and holes were drilled (coordinates: PVN [AP=-0.8, ML=±0.2, DV=-5.5], lateral ventricle for cannula implantation [AP = 0.3, ML = 1.0, D = 2.5]). The AAVs were injected at a rate of 0.1μL/min, and guide cannulas (PlasticsOne, 2.5mm) were implanted and secured to the skull using a tissue bonding glue (Loctite 454) and dental acrylic, and closed with a screw-on dummy cannula. Animals were given preoperative analgesic (buprenorphine, 0.3 mg/kg) and post-operative anti-inflammatory Meloxicam (5mg/Kg) and allowed to recover at least 10 days during which time they were singly-housed and handled frequently.

### Mouse metabolism studies

Mice were single housed after AAV injection. Weight was measured weekly, or as mentioned in figures. Fat mass and lean mass were measured using EchoMRI (EchoMRI LLC, Houston, TX). Food intake was assessed either by hand or measured by CLAMS (Columbus Instruments, Colombus, OH). Energy expenditure was measured by CLAMS (Columbus Instruments, Colombus, OH). Mice were tested over 96 continuous hours, and the data from the last 48 hours were analyzed. Energy expenditure (EE) is expressed in terms of kcal per hour and calculated using the Lusk equation: EE=(3.815+1.232×RER)×VO2.

| Figure | EchoMRI | Food intake | Energy Expenditure |
| --- | --- | --- | --- |
| Figure 1 | Tx+5, +12, +22, +27days | By CLAMS: Tx+6-9, +13-16; +23-36 days | By CLAMS, Tx+6-9, T13-16, +23-36 days |
| Figure 2 | / | Post aCSF/MTII ICV injection food intake: 4h- food intake manually measured following a 24h fast at Tx-2, +2, +11, +18 days, prior to weight divergence | / |
| Figure 3 | / | Post aCSF/MTII ICV injection food intake: 4h-food intake manually measured following a 24h fast. | / |
| Figure 4 | At 12 weeks old | 24h food intake measured manually daily over 4 days | / |
| Figure 5 | AAV +0, 3, 6 and 9 weeks | By CLAMS at AAV+ 3 weeks | By CLAMS at AAV+ 3 weeks |

### Direct fluorescence and immunofluorescence studies of mouse hypothalamus

Mice were perfused trans-cardially with PBS followed by 4% paraformaldehyde fixation solution. Brains were dissected and post-fixed in fixation solution at 4°C overnight, soaked in 30% sucrose solution overnight, embedded in O.C.T. (Tissue-Tek, Sakura Finetek USA, INC., Torrance, CA), frozen, and cut into 20-35 μm coronal sections, then stored at −80°C until staining. After washing, sections were blocked for 1 hr in 50% serum 50% antibody buffer (1.125, %NaCL, 0.75%Tris base, 1%BSA,1.8 %L-Lysine, 0.04% azide), followed by incubation with primary antibody overnight at 4°C. After washing, sections were incubated with secondary antibodies for an hour at room temperature, washed and stained with Hoechst (1:5000), washed and mounted with Prolong™ Diamond antifade Mountant.

| Primary antibodies |  |
| --- | --- |
| Chicken anti-GFP (abcam, ab13970) | 1:250 |
| Rabbit anti-Adcy3 (Santa Cruz Biotechnology, sc-588) | 1:500 |
| Mouse anti-Flag M2 (Sigma F1804) | 1:500 |
| Rabbit anti-Ps6 ribosomal protein (Invitrogen, 44-923-G) | 1:1000 |
| Secondary antibodies |  |
| Goat anti-chicken Alexa fluor 488 (Invitrogen) | 1:500 |
| Goat anti-rabbit Alexa fluor 633 or 555 or 488 (Invitrogen) | 1:500 |
| Goat anti-mouse Alexa fluor 555 (Invitrogen) | 1:500 |

### Image capture and processing

Confocal images were generated using a Leica SP5 (Supplementary Figure 1, 3), a Zeiss LSM 780 confocal microscope (Figure 3) or a Nikon W1 spinning disk confocal (Figure 5). Images were processed with Fiji. Maximal intensity Z projections are from at least 20 slices over 10-20 μm. Widefield Images of Ps6 staining (Supplementary Figure 3) were generated using an Zeiss Apotome.

### Tamoxifen-induced recombination

*Ubc-Cre-Ert2-dependent* recombination was induced by intra-peritoneal administration of tamoxifen (Sigma-Aldrich, St. Louis, MO) for 5 consecutive days at a dose of 10.0mg/ 40g body weight.

### Anorexigenic response to MTII

Experiments were performed as described^36^. Mice were deprived of food for 24h and injected intracerebroventricularly with either 1 μl artificial cerebro-spinal fluid (aCSF) or MTII (Genescript, Piscataway, NJ, 0.5nmoles in experiment in Figure 3 and 0.05nmoles for experiment in Figure 4) in 1μl aCSF over a 15 sec period. Each mouse was returned to its home cage and a weighed amount of chow was placed into the cage. The remaining chow was weighed after 4 hours to determine intake. Permeability and placement of the cannula was assessed before and after the protocol by measuring the drinking response to angiotensin II.

### Anorexigenic response to THIQ

Experiments were performed as described^37^. Mice were deprived of food at 5PM, injected with 1 μl aCSF or 1μl (32nM) THIQ (Tocris Bioscience, Bristol, UK) in aCSF at 7PM and food intake was measured 4 hours later.

### Osmotic stimulation

Mice were given an IP injection of 2 M NaCl (350 μl) or normal saline (0.9% NaCl, 350 μl) as control, water was removed from the cage, and mice were allowed access to food and were perfused 120 min later.

### Hypothalamic RNA and Protein quantification

Hypothalamic tissue was collected by dissection. Ift88 protein levels were assayed by immunoblot (as previously described by Reed et al. 2010).

| Primary antibody | Secondary antibody |
| --- | --- |
| Rabbit anti-Ift88 (Proteintech, 13967-1-AP), 1:500 | Goat anti-rabbit IgG-HRP (Santa Cruz) 1:3000 |
| Mouse anti-GAPDH (6C5) (Santa Cruz Biotechnology, 1:1000) | Goat anti-mouse IgG-HRP (Santa Cruz) 1:3000 |

mRNA levels for *Agrp, Npy, Pomc, Mc4r, Sim1, Ift88, β-actin* were assayed by quantitative RT-PCR using Taqman assays.

### Statistical analysis

Sample sizes were chosen based upon the estimated effect size drawn from previous publications (Siljee et al., 2018) and from the performed experiments. Data distribution was assumed to be normal, but this was not formally tested. Statistical analysis was performed using unpaired Student t test or repeated measures of two-way ANOVA followed by Sidak’s multiple comparisons test, as indicated. All data were expressed as mean ± SEM. A p-value ≤0.05 was considered as statistically significant. All data were analyzed using Prism 8.0 (GraphPad Software).

