## Supplemental data for "The primary cilium is required for MC4R control of food intake and body weight"

### SUPPLEMENTAL FIGURES:

A

| Summary of changes in *Ubc-Cre-Ert2 Ift88$^{fl/fl}$* mice after Tamoxifen Injection | Tx+7d | Tx+15d | Tx+22d |
|---|---|---|---|
| mRNA Expression (% of control) | 9.1 | 6.3 | 6.4 |
| Protein Expression (% of control) | 50.0 | 22.5 | 17.7 |
| Relative ADCY3 IF | +++ | + | +/- |

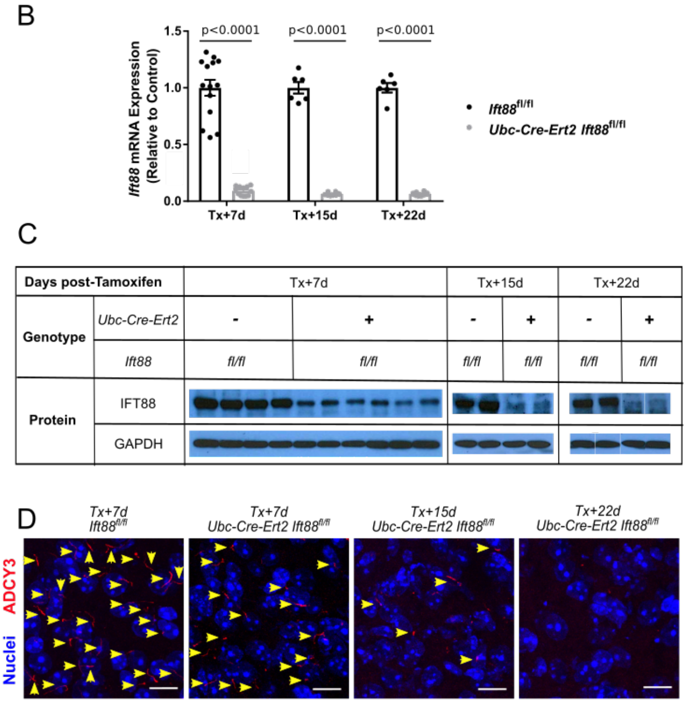


**Supplementary Figure 1: Time course of IFT88 loss and cilia ablation following tamoxifen injection in *UBC-cre/ERT2 Ift88$^{fl/fl}$* mice. A)** Summary of time dependent changes in mRNA expression, protein expression and cilia ablation. **B)** Time–course of hypothalamic *Ift88* mRNA expression measured by qPCR following tamoxifen injection in *Ift88$^{fl/fl}$* and *UBC-cre/ERT2 Ift88$^{fl/fl}$* mice. **C)** Time course of hypothalamic IFT88 protein expression measured by western blots following tamoxifen injection in *Ift88$^{fl/fl}$* and *UBC-cre/ERT2 Ift88$^{fl/fl}$* mice. **D)** Immunofluorescence of ADCY3 (red), a neuronal primary cilia specific protein, allows for the time course of cilia loss (arrows) in the hypothalamus of *UBC-cre/ERT2 Ift88*$^{fl/fl}$ mice following tamoxifen injection.

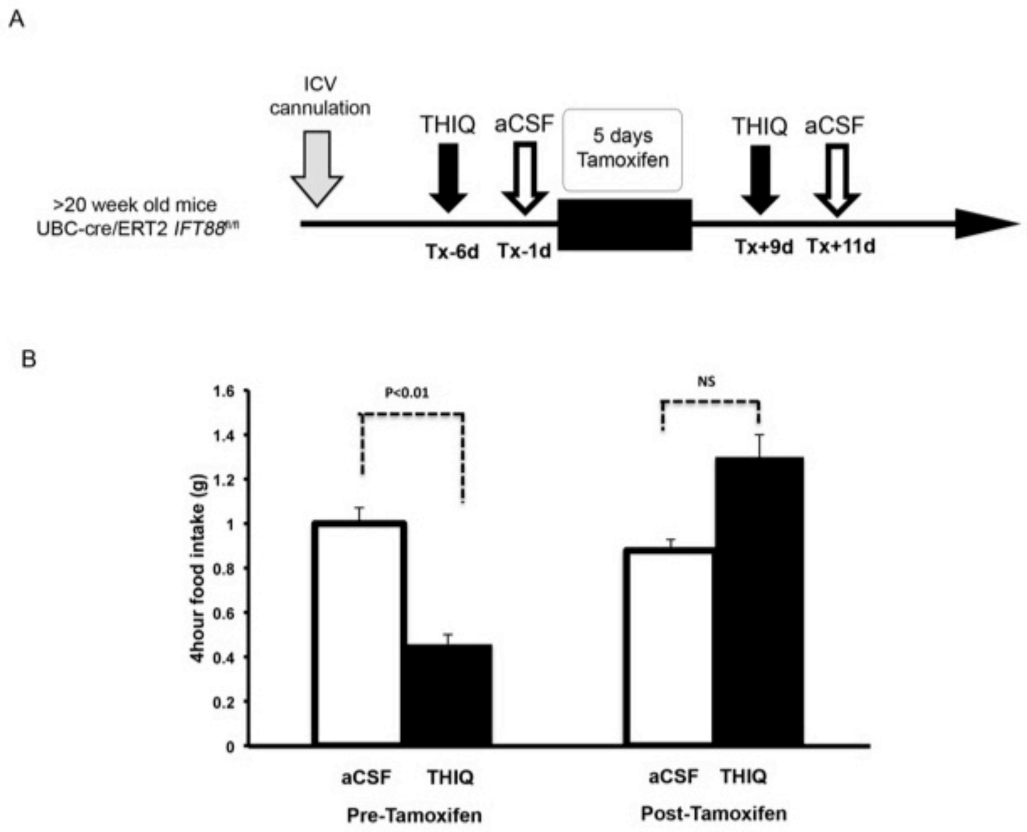


**Supplementary Figure 2: Loss of the primary cilium abolishes THIQ activation of MC4R. A)** Experimental protocol. **B)** Loss of the anorectic effects of ICV THIQ in tamoxifen treated *UBC-cre/ERT2 Ift88*$^{fl/fl}$ mice. Data are represented as mean+/-SEM.

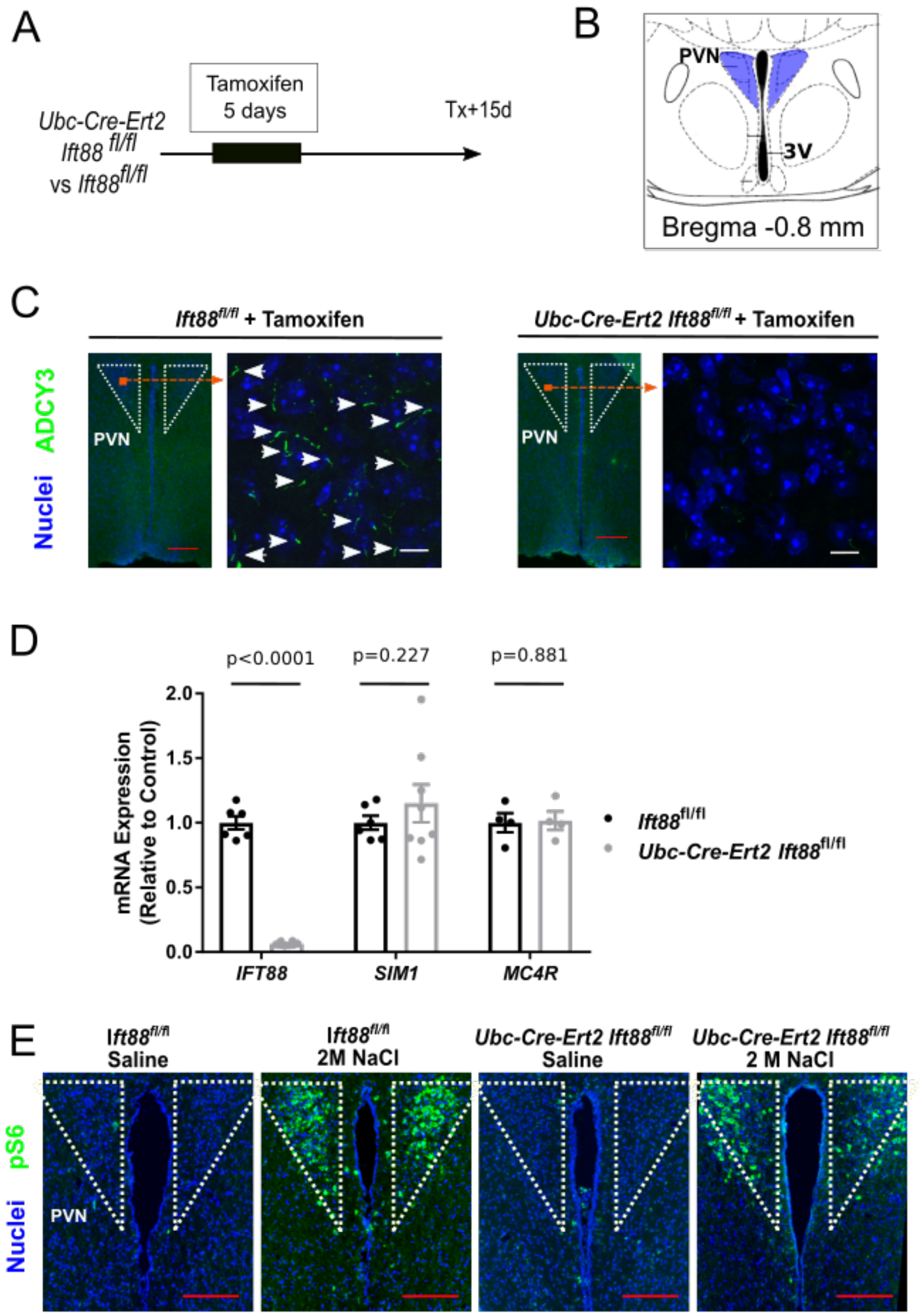


**Supplementary Figure 3: Complete loss of PVN cilia does not affect hypothalamic MC4R expression or general PVN neuronal function. A)** Experimental protocol: 20 week old mice were treated with tamoxifen and their brains were harvested 15 days later. **B)** Schematic representation of hypothalamic region studied **C)** ADCY3-labeled primary cilia (green, arrows) are detected in neurons of the PVN of $Ift88^{fl/fl}$ but not *UBC-cre/ERT2* $Ift88^{fl/fl}$ mice. **D)** *Sim1* and *MC4R* mRNA levels are unaffected in *UBC-cre/ERT2* $Ift88^{fl/fl}$ (n=4) vs $Ift88^{fl/fl}$ (n=3) male mice at 15 days following tamoxifen administration. **E)** Immunofluorescence staining of phospho-S6 in $Ift88^{fl/fl}$ and *UBC-cre/ERT2* $Ift88^{fl/fl}$ mice, 15 days after tamoxifen injection and 2 hours after either 0.3M (saline) or 2 M NaCl injection. Red scale bars represent 200 µm, white scale bar represent 20 µm. PVN: paraventricular nucleus of the hypothalamus; 3V: third ventricle.
